# In-shoe plantar pressure depends on walking speed and type of weight-bearing activity in people with diabetes at high risk of ulceration

**DOI:** 10.1101/2022.02.09.479511

**Authors:** Chantal M Hulshof, Jaap J van Netten, Maartje G Dekker, Mirjam Pijnappels, Sicco A Bus

## Abstract

**Background:** In evaluating the biomechanical properties of therapeutic footwear, most often in-shoe plantar pressure is obtained during mid-gait steps at self-selected speed in a laboratory setting. However, this may not represent plantar pressures or indicate the cumulative stress experienced in daily life, where people adopt different walking speeds and weight-bearing activities.

**Research question:** In people with diabetes at high risk of ulceration, 1) what is the effect of walking speed on plantar pressure measures, and 2) what is the difference in plantar pressure measures between walking at self-selected speed and other weight-bearing activities?

**Methods:** In a cross-sectional study, we included 59 feet of 30 participants (5 females, mean age: 63.8 (SD 9.2) years). We assessed in-shoe plantar pressure with the Pedar-X system during three standardized walking speeds (0.8, 0.6 and 0.4 m/s) and eight types of activities versus walking at self-selected speed (3 components of the Timed Up and Go test (TUG), standing, accelerating, decelerating, stair ascending and descending and standing). Peak plantar pressure (PPP) and pressure-time integral (PTI) were determined for the hallux, metatarsal 1, metatarsal 2-3 and metatarsal 4-5. For statistical comparisons we used linear mixed models (α<0.05) with Holm-Bonferroni correction.

**Results:** With increasing walking speed, PPP increased and PTI decreased for all regions (p≤0.001). Standing, decelerating, stair ascending and TUG showed lower PPP than walking at self-selected speed for most regions (p≤0.004), whereas accelerating and stair descending showed similar PPP. Stair ascending and descending showed higher PTI than walking at self-selected speed (p≤0.002), standing showed lower PTI (p≤0.001), while the other activities showed similar PTI for most regions.

**Significance:** To best evaluate the biomechanical properties of therapeutic footwear, and to assess cumulative plantar tissue stress of people with diabetes at high risk of ulceration, plantar pressures during different walking speeds and activities of daily living should be considered.

## 1. Introduction

High mechanical loading is a frequent cause of diabetes-related foot ulcers [1]. High mechanical loading results from changes in foot structure and sensation that occur when longstanding diabetes mellitus leads to peripheral neuropathy [2]. Of people with diabetes, 19-34% develop a foot ulcer in their life [3]. With the current 451 million people with diabetes, and this number rising, the burden of diabetes-related foot ulcers is large, and will increase further [4]. By reducing mechanical load on the foot, foot ulcer incidence can be reduced, reducing the burden of diabetes-related foot ulcers.

Mechanical loading of the foot in diabetes is mostly expressed by the plantar pressure distribution [1]. By redistributing pressure from high pressure locations to low pressure locations on the plantar side of the foot, high plantar pressure locations can be offloaded. Such offloading can be achieved with therapeutic footwear, as recommended in international guidelines [5]. Therapeutic footwear with adequate offloading can help prevent foot ulcers, provided they are worn. However, even in optimized therapeutic footwear with proven offloading effect, diabetes-related foot ulcers develop [6].

The offloading effect of therapeutic footwear is typically evaluated by measuring in-shoe plantar pressure during mid-gait steps while walking overground at self-selected speed in a laboratory setting [1]. However, this may not represent the plantar pressure or the cumulative plantar tissue stress experienced in daily life [7]. In a laboratory setting, people are assessed in optimal circumstances: unperturbed and undistracted walking in a straight line over a flat surface at their self-selected speed, most often their comfortable speed. Studies show that the comfortable walking speed of people in a laboratory setting is faster than the most common walking speed in daily life [8,9]. Furthermore, in their daily life setting, people vary in weight-bearing activities with, for example, standing, turning, accelerating, decelerating and stair walking in addition to overground walking at a constant speed. Such variety in speed and activities may affect the cumulative plantar tissue stress and the evaluation of treatments for prevention and healing of diabetes-related foot ulceration.

The effect of walking speed on plantar pressure has only been investigated in healthy people [10–15]. These studies show that increased walking speed results in increased peak plantar pressure (PPP) and decreased pressure-time integral (PTI) in most foot regions [10–15]. However, only one of these studies measured pressures during overground walking, all others used a treadmill, which may affect pressures measured [16]. The one study that measured plantar pressure during overground walking used 0.95, 1.33 and 1.62 m/s as controlled walking speeds [12]. However, people with diabetes at high risk of ulceration generally walk slower [17], with smaller expected variation in walking speed in daily life.

Plantar pressure during different activities of daily living has been investigated in healthy people [18– 20], people with diabetes [21], and in people with diabetes and neuropathy [22,23], but not in those at high risk of ulceration. These studies showed that in-shoe PPP was lower during stair walking, the Timed Up and Go test (TUG), and ramp ascending and descending, compared to overground walking [18–23]. In people with diabetes, in-shoe PPP was lower during turning compared to overground walking [21,22], while the opposite was shown in healthy people [18]. In healthy people and people with diabetes, PTI was higher in the metatarsal region during stair walking than during overground walking [20,22], while no difference was found for turning and ramp ascending and descending [22]. While these studies provide some insights, plantar pressures were not investigated during other typical weight-bearing activities of daily living, such as standing, and accelerating or decelerating walking steps. People with diabetes at high ulcer risk spend twice as much time standing than walking [24], and have on average more than 350 bouts of walking per day [24], with each bout having accelerating and decelerating steps, justifying the investigation of more activities of daily living in this patient group.

More insight into the in-shoe plantar pressure distribution of people with diabetes at high risk of ulceration in a variety of speed and activity conditions can help in building better biomechanical models of diabetic foot ulceration and prevention, and can help to improve the assessment of the offloading effect of therapeutic footwear. We therefore aimed to 1) investigate the effect of walking speed on plantar pressure measures, and 2) compare plantar pressure measures of overground walking at self-selected speed with plantar pressure measured during different weight-bearing activities of daily living, in people with diabetes at high risk of ulceration.

## 2. Methods

### 2.1. Participants

In a cross-sectional study design, we included 59 feet of 30 ambulant participants of 18 years and older with diabetes mellitus, loss of protective sensation, a recently healed foot ulcer (<1 year) or high barefoot plantar pressures (>600 kPa at any region in either foot). All particpants were stratified as IWGDF risk 3 [2]. One person had only one foot, following a transtibial amputation seven months before inclusion.

Exclusion criteria were a foot ulcer, open amputation wound, active Charcot neuro-osteo arthropathy, use of a walking aid for full support or critical ischemia (toe pressure <30 mmHg [25]). This study is part of an ongoing multi-center prospective observational cohort study registered in the Netherlands trial register (registration number: NL8839). Written informed consent was obtained from all participants prior to inclusion. All study procedures were in accordance with the Declaration of Helsinki. The requirement for ethical review of the study was waived under the Medical Research Involving Human Subjects Act in the Netherlands by the local medical ethics committee of Amsterdam UMC (registration number: W19_429#19.495).

### 2.2. Measurements and equipment

During clinical examination we examined participants’ feet for deformities, bony prominences, amputations and pre-signs of ulceration, and we screened participants for their medical and ulcer history. Subsequently, we assessed loss of protective sensation using a 10-gram monofilament and tuning fork, and vascular status by testing pedal pulses and measuring toe pressures, both according to international guidelines [2]. Barefoot dynamic plantar pressure during walking was measured at a frequency of 100 Hz with the EMED-X platform with 4 sensors/cm^2^ (Novel GmbH, Munich, Germany), using the 2-step gait approach protocol [26]. Mean PPP for the forefoot was calculated over four steps.

In-shoe plantar pressures were obtained during three speed and nine activity conditions (Table 1). The order of tested trials was: 1) standing followed by walking at self-selected speed, including accelerating and decelerating at the start and end of the trial, 2) walking at the three standardized speeds, 3) Timed Up and Go test (TUG) and 4) stair walking. To measure in-shoe plantar pressures, we used Pedar-X (Novel GmbH, Munich, Germany), consisting of calibrated 2 mm thick flexible insoles with 99 capacitive sensors (∼1 sensor/2 cm^2^), measures at 50 Hz and with high accuracy and repeatability [27]. Six different sizes of wide insoles were available to accommodate a range of shoe sizes. All in-shoe plantar pressures were measured at the sock-insole interface in the participant’s most frequently used pair of shoes. Walking speed was determined during steady-state walking using a photocell system (Tag Heuer, Neuchâtel, Swiss) located along a 12m-walkway. For the standardized walking speed trials, light-emitting diodes were placed on the ground projecting a fixed speed moving pattern to be followed by the participants. The TUG and its conditions and the stair walking trials were timed with a MoveTest intertial sensor (McRoberts, Den Haag, the Netherlands), containing 3D accelerometers and gyroscopes, worn dorsally at vertebra L5 using an elastic belt.

**Table 1.** Detailed overview of all test conditions

|  | Trial<sup>a</sup> | Condition | Instructions for the participant | Number of trials | Location | Time registration |
| --- | --- | --- | --- | --- | --- | --- |
| **Speed conditions** | **Walking at standardized walking speeds<sup>b</sup>** | 1. 0.8 m/s<br>2. 0.6 m/s<br>3. 0.4 m/s | Follow the moving LEDs on the floor. Before the first attempt at each speed, a test trial was performed for familiarization. | 2-4 trials (depending on number of steps per trial, $\geq 12$ midgait steps needed [30]) | 6m LED strip along the 12-m walkway in the gait laboratory | Photocell system<br>- 5% inter-trial variation allowed |
| **Activity conditions** | **Walking at self-selected speed** | 1. Standing<br>2. Accelerating<br>3. Walking<br>4. Decelerating | Stand still in your natural position for four seconds, walk at comfortable speed to the other side of the laboratory and stand still in your natural position again for four seconds. | 2-4 trials (depending on number of steps per trial, $\geq 12$ midgait steps needed [30]) | 12m-walkway in the gait laboratory | Photocell system<br>- 10% inter-trial variation allowed |
|  | **Timed Up and Go test (TUG)** | 5. Sit-to-stand<br>6. Walking including turning<br>7. Stand-to-sit | Stand up from a chair, walk to the cone (3 meters distance), turn around the cone, walk back to the chair and sit down, all at comfortable speed [39]. | 2 clockwise<br>2 counter-clockwise | TUG setup in the gait laboratory | MoveTest |
| | **Stair walking** | 8. Ascending<br>9. Descending | Stand still close to the staircase, ascend and descend the staircase in a comfortable way and stand still after completing stair walking. Use of handrails was allowed, available at both sides of the staircase. | 3 ascending<br>3 descending<br>($\geq 10$ stair steps needed [40]) | Staircase with 9-steps (step height: 18.5 cm, step depth: 27 cm, 34° slope) directly outside the gait laboratory, unperturbed by other people | MoveTest |

<sup>a</sup> = The order of trials was: 1) walking at self-selected speed, 2) walking at standardized walking speeds, 3) Timed Up and Go test and 4) stair walking. <sup>b</sup> = The three speeds were 0.8, 0.6 and 0.4 m/s, selected as range of most common walking speeds during daily life activities [8]. (LED = light-emitting diodes)

### 2.3. Outcome measures

To decide on using the foot or the person as level of analysis [28], we calculated the pearson correlation coefficient between the barefoot plantar pressures of the left and right forefoot; this was 0.437, a low-to-moderate coefficient. We therefore considered the data of the left and right foot as independent for outcomes on PPP and PTI, and all analyses were done on foot level.

PPP and PTI were calculated for each condition at four forefoot regions: the hallux, metatarsal 1, metatarsal 2-3 and metatarsal 4-5, those being the most susceptible locations for foot ulceration [29]. PPP was defined as the highest pressure measured during stance. PTI was defined as the area under the peak pressure-time curve during stance, determined by trapezoidal numerical integration. PTI is time-dependent and because standing does not contain a step, we used one second of standing time to determine PTI, comparable to the stance time in walking. Both PPP and PTI were averaged over all valid steps recorded. We extracted raw force and pressure data from the Novel software (Novel GmbH, Munich, Germany). Mid-gait steps, accelerating and decelerating, standing, the TUG components and stair steps were detected based on the characteristics of the force data, using custom-made Matlab scripts (Mathworks, California, United States, version R2021a). If a participant’s self-selected walking speed was lower than 0.8 m/s, this participant only walked at the standardized speeds of 0.4 and 0.6 m/s, and plantar pressure at 0.8 m/s was based on the mean normalized change in plantar pressure from 0.6 to 0.8 m/s of all other participants.

### 2.4. Statistical analysis

All analyses were performed with PPP and PTI as outcome measures, using SPSS version 26. We used a convenience sample of 30 participants [30]. Normality of PPP and PTI data were checked with the Shapiro-Wilk test.

To analyze the effect of walking speed on plantar pressure, we used linear mixed model analysis of variance (ANOVA) with the forefoot regions (n=4) as between factor and the speed conditions (n=3) as within factor. With a total 12 comparisons, we used the Holm-Bonferroni method to correct for multiple testing, with alpha 0.05. This correction means that we sorted the p-values from lowest to highest, and compared the resulting p-values to corrected nominal alpha levels with alpha=alpha/12=0.05/12=0.004167 for the smallest p-value; alpha/11 for the second smallest, etc. When the p-value was smaller than its corrected nominal alpha value, a result was considered statistically significant.

To compare plantar pressure during walking at self-selected speed with the other activities of daily living, we used one repeated measures anova per foot region (n=4) with activity conditions (n=9) as within factor. We only compared walking at self-selected speed with each activity condition separately, resulting in eight comparisons per foot region, with alpha at 0.05 and the Holm-Bonferroni method to correct for multiple testing (alpha=alpha/8=0.05/8=0.00625 for the smallest p-value; etc.).

## 3. Results

Baseline descriptive characteristics and outcome data of the 30 included participants and their 59 feet are shown in Table 2.

**Table 2.** Baseline descriptive characteristics and outcome data

|  |  |
| --- | --- |
| **Participants** | **N=30** |
| **Age (years)** | 63.8±9.2 |
| **Sex (female/male)** | 16.7% (5) / 83.3% (25) |
| **Diabetes type (1/2)** | 30.0% (9) / 70.0% (21) |
| **Diabetes duration (years)** | 18.0±11.5 |
| **BMI (kg/m<sup>2</sup>)** | 30.0±5.5 |
| **HbA1c (mmol/mol)** | 61.6±18.6 |
| **Loss of sensation** |  |
| - Loss of both pressure and vibratory sensation | 90.0% (27) |
| - Loss of only pressure sensation | 10.0% (3) |
| - Loss of only vibratory sensation | 0.0% (0) |
| **Ulcer history** | 96.7% (29) |
| - Recent (<1 year) | 50.0% (15) |
| **Time since healing of last ulcer (months)** | 23.4±46.6 |
| **Foot (most recent ulcer)** |  |
| - Left | 48.3% (14)<sup>a</sup> |
| - Right | 51.7% (15)<sup>a</sup> |
| **Ulcer location (most recent ulcer)** |  |
| Plantar | 69.0% (20)<sup>a</sup> |
| - Hallux | 24.1% (7)<sup>a</sup> |
| - Digit 4 | 13.8% (4)<sup>a</sup> |
| - MTP 1 | 6.9% (2)<sup>a</sup> |
| - MTP 2 | 3.4% (1)<sup>a</sup> |
| - MTP 5 | 10.3% (3)<sup>a</sup> |
| - Medial midfoot | 6.9% (2)<sup>a</sup> |
| - Calcaneus | 3.4% (1)<sup>a</sup> |
| Dorsal | 27.5% (8)<sup>a</sup> |
| - Hallux | 10.3% (3)<sup>a</sup> |
| - Digit 2 | 3.4% (1)<sup>a</sup> |
| - Digit 3 | 10.3% (3)<sup>a</sup> |
| - Digit 4 | 3.4% (1)<sup>a</sup> |
| Unknown | 3.4% (1)<sup>a</sup> |
| **Walking speed (m/s)** |  |
| - Walking at self-selected speed<sup>b</sup> | 1.1±0.2 |
| - TUG walking | 0.9±0.3 |
| - Stair ascending | 0.5±0.1 |
| - Stair descending | 0.5±0.1 |
| **TUG duration (s)** | 12.7±2.8 |
| - Sit-to-stand | 2.0±0.5 |
| - Walking including turning | 8.2±2.4 |
| - Stand-to-sit | 2.5±0.5 |
| **Step duration (s)** |  |
| - Walking at self-selected speed | 0.9±0.1 |
| - TUG walking | 1.1±0.1 |
| - Stair ascending | 1.1±0.3 |
| - Stair descending | 1.1±0.3 |
| - 0.4 m/s | 1.5±0.2 |
| - 0.6 m/s | 1.2±0.2 |
| - 0.8 m/s | 1.1±0.1 |
| **Feet** | **N=59** |
| **Barefoot PPP (kPa) during walking<sup>c</sup>** |  |
| - Left foot | 902.2±259.3 |
| - Right foot | 929.2±276.1 |

| Foot deformities<sup>d</sup> |  |
| --- | --- |
| - Absent | 1.7% (1) |
| - Mild | 6.8% (4) |
| - Moderate | 78.0% (46) |
| - Severe | 13.6% (8) |
| Amputations |  |
| - Digits 1-5 | 22.0% (13) |
| - Metatarsal region | 3.4% (2) |
| - No amputation | 74.6% (44) |

Continuous data are mean ± standard deviation and discrete data are percentage (number). <sup>a</sup> = percentage relative to people with ulcer history ( $n=29$ ), <sup>b</sup> = difference in walking speed between walking at self-selected speed and TUG walking including turning is 0.2 m/s. <sup>c</sup> = 29 participants had barefoot plantar pressure >600 kPa. <sup>d</sup> = foot deformity was classified as absent, mild, moderate and severe [6]. (BMI = body mass index, HbA1c = glycated hemoglobin, TUG = Timed Up and Go test, PPP = peak plantar pressure)

For the standardized walking speeds, in 11 of the 12 comparisons of speed and regions, PPP increased significantly with increased walking speed (p<0.001) (Figure 1, Table 3). PTI decreased significantly with increased walking speed for all walking speeds and regions (p≤0.010).

**Figure 1:**
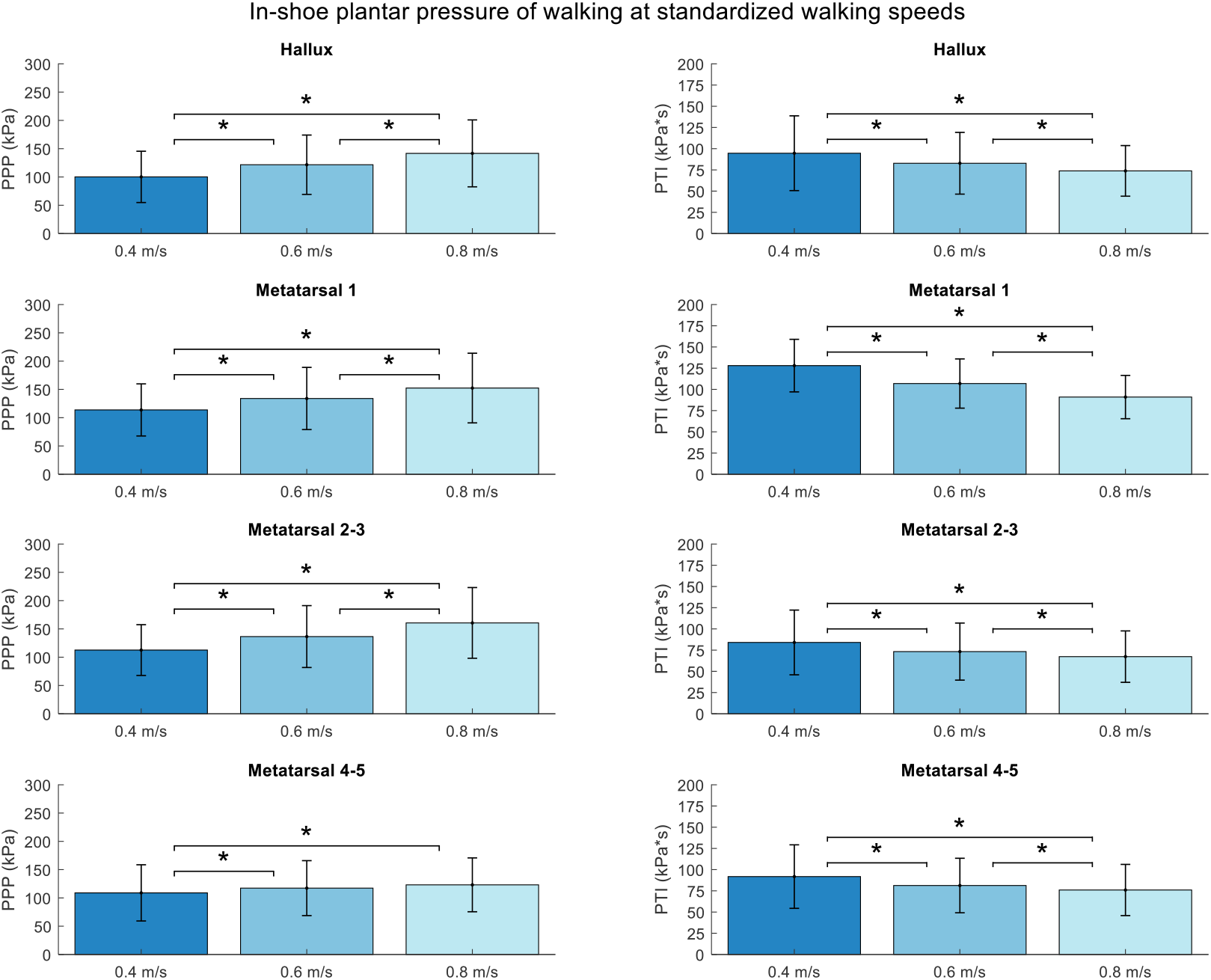
Peak pressure (PP) and pressure-time integral (PTI) (mean ± standard deviation) of standardized walking speeds. * = significantly different from each other; cut-off for p-values to be considered statistically significant was corrected for multiple testing with the Holm-Bonferroni method.

**Table 3.**
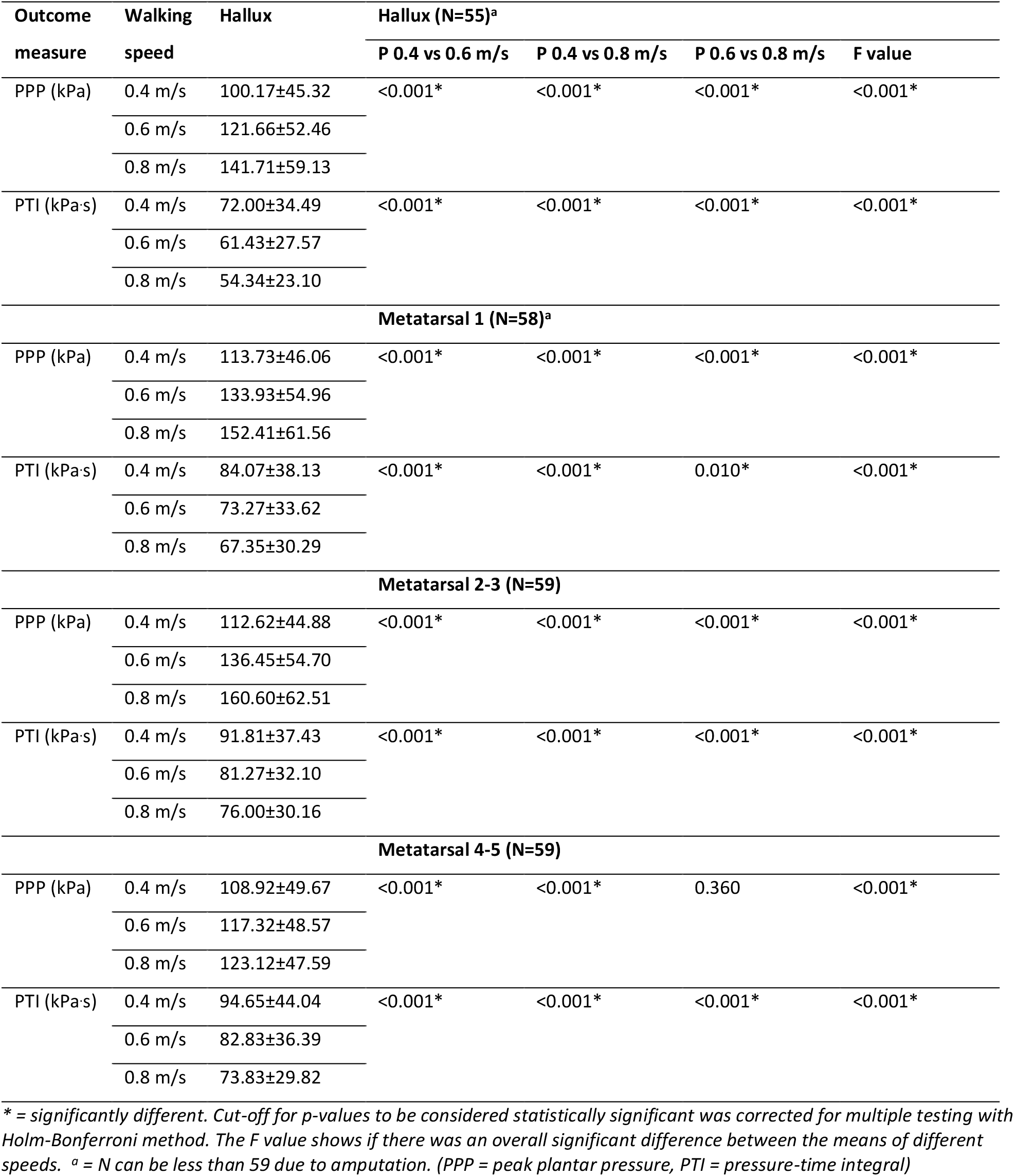
Detailed results of plantar pressure measures during standardized walking speeds

| Outcome measure | Walking speed | Hallux | Hallux (N=55)<sup>a</sup> |  |  |  |
| --- | --- | --- | --- | --- | --- | --- |
|  |  |  | P 0.4 vs 0.6 m/s | P 0.4 vs 0.8 m/s | P 0.6 vs 0.8 m/s | F value |
| PPP (kPa) | 0.4 m/s | 100.17±45.32 | <0.001* | <0.001* | <0.001* | <0.001* |
|  | 0.6 m/s | 121.66±52.46 |  |  |  |  |
|  | 0.8 m/s | 141.71±59.13 |  |  |  |  |
| PTI (kPa-s) | 0.4 m/s | 72.00±34.49 | <0.001* | <0.001* | <0.001* | <0.001* |
|  | 0.6 m/s | 61.43±27.57 |  |  |  |  |
|  | 0.8 m/s | 54.34±23.10 |  |  |  |  |
| Metatarsal 1 (N=58)<sup>a</sup> |  |  |  |  |  |  |
| PPP (kPa) | 0.4 m/s | 113.73±46.06 | <0.001* | <0.001* | <0.001* | <0.001* |
|  | 0.6 m/s | 133.93±54.96 |  |  |  |  |
|  | 0.8 m/s | 152.41±61.56 |  |  |  |  |
| PTI (kPa-s) | 0.4 m/s | 84.07±38.13 | <0.001* | <0.001* | 0.010* | <0.001* |
|  | 0.6 m/s | 73.27±33.62 |  |  |  |  |
|  | 0.8 m/s | 67.35±30.29 |  |  |  |  |
| Metatarsal 2-3 (N=59) |  |  |  |  |  |  |
| PPP (kPa) | 0.4 m/s | 112.62±44.88 | <0.001* | <0.001* | <0.001* | <0.001* |
|  | 0.6 m/s | 136.45±54.70 |  |  |  |  |
|  | 0.8 m/s | 160.60±62.51 |  |  |  |  |
| PTI (kPa-s) | 0.4 m/s | 91.81±37.43 | <0.001* | <0.001* | <0.001* | <0.001* |
|  | 0.6 m/s | 81.27±32.10 |  |  |  |  |
|  | 0.8 m/s | 76.00±30.16 |  |  |  |  |
| Metatarsal 4-5 (N=59) |  |  |  |  |  |  |
| PPP (kPa) | 0.4 m/s | 108.92±49.67 | <0.001* | <0.001* | 0.360 | <0.001* |
|  | 0.6 m/s | 117.32±48.57 |  |  |  |  |
|  | 0.8 m/s | 123.12±47.59 |  |  |  |  |
| PTI (kPa-s) | 0.4 m/s | 94.65±44.04 | <0.001* | <0.001* | <0.001* | <0.001* |
|  | 0.6 m/s | 82.83±36.39 |  |  |  |  |
|  | 0.8 m/s | 73.83±29.82 |  |  |  |  |

\* = significantly different. Cut-off for p-values to be considered statistically significant was corrected for multiple testing with Holm-Bonferroni method. The F value shows if there was an overall significant difference between the means of different speeds. <sup>a</sup> = N can be less than 59 due to amputation. (PPP = peak plantar pressure, PTI = pressure-time integral)

Overall, all TUG components, decelerating, stair ascending and standing showed signifcantly lower PPP than walking at self-selected speed for all regions (p≤0.004), while PPP was similar for accelerating and stair descending compared to walking at self-selected speed for all regions (p≥0.029) (Figure 2, Table 4). In general, PTI was significantly higher for stair ascending and descending (p≤0.002), significantly lower for standing (p≤0.001), and similar for all TUG components, accelerating and decelerating for all regions (p≥0.043), compared to walking at self-selected speed.

**Figure 2:**
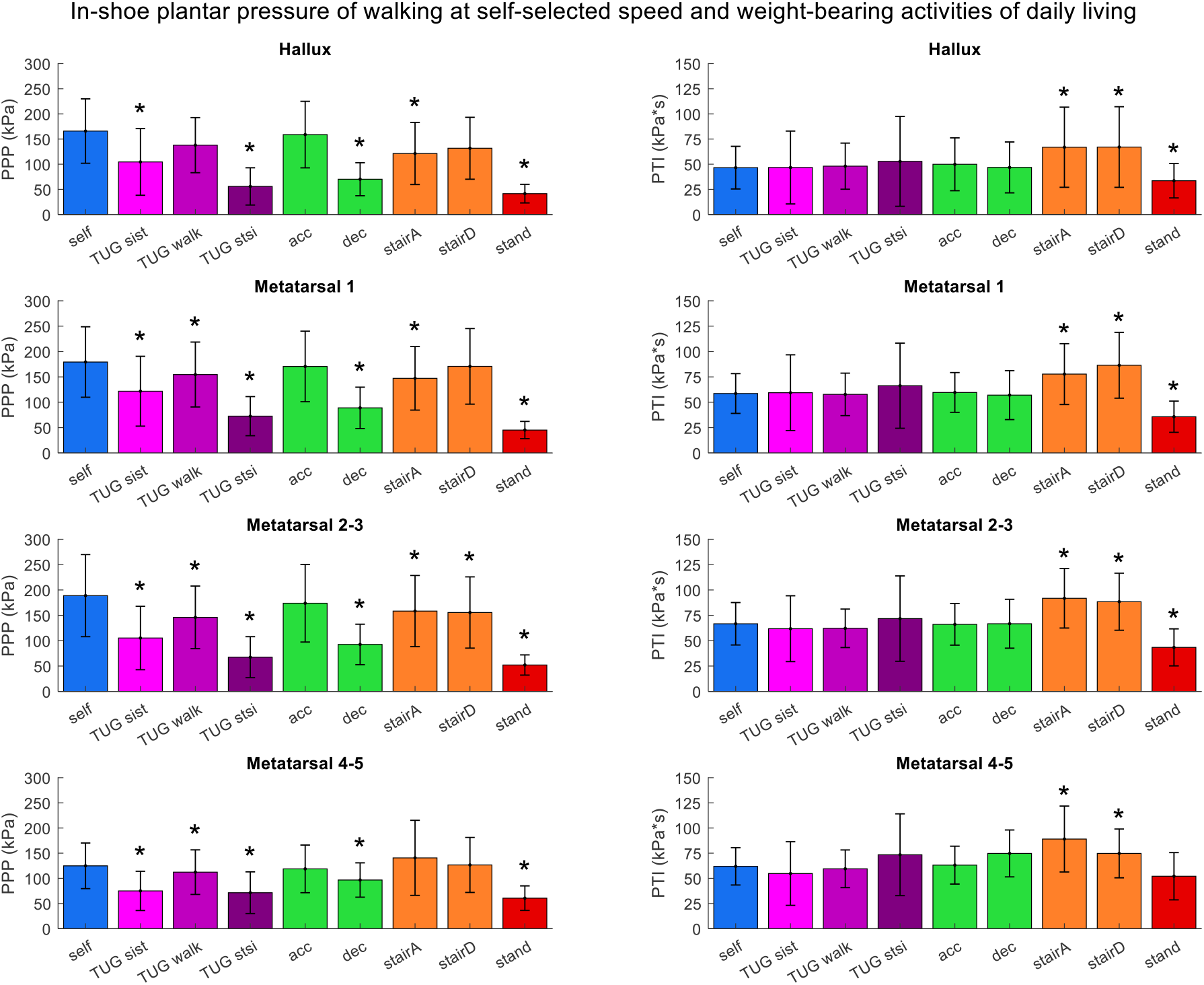
Peak pressure (PP) and pressure-time integral (PTI) (mean ± standard deviation) during different activities of daily living. * = significantly different from walking at self-selected speed. Cut-off for p-values to be considered statistically significant was corrected for multiple testing with Holm-Bonferroni method. (self = walking at self-selected speed, TUG = Timed Up and Go test,; sist = sit-to-stand; walk = walking including turning; stsi = stand-to-sit;, acc = accelerating, dec = decelerating, stairA = stair ascending, stairD = stair descending, stand = standing)

**Table 4.** Detailed results of plantar pressure measures during activities of daily living

| Outcome measure | Activity | Hallux (N=55)<sup>a</sup> |  |  | Metatarsal 1 (N=58)<sup>a</sup> |  |  | Metatarsal 2-3 (N=59) |  |  | Metatarsal 4-5 (N=59) |  |  |
| --- | --- | --- | --- | --- | --- | --- | --- | --- | --- | --- | --- | --- | --- |
|  |  | Mean pressure | P value | F value | Mean pressure | P value | F value | Mean pressure | P value | F value | Mean pressure | P value | F value |
| PPP (kPa) | self | 165.86±63.97 | - | <0.001* | 179.34±69.45 | - | <0.001* | 188.85±80.92 | - | <0.001* | 124.87±45.30 | - | <0.001* |
|  | TUGsist | 104.53±66.27 | <0.001* |  | 121.79±68.83 | <0.001* |  | 105.28±62.49 | <0.001* |  | 75.02±39.01 | <0.001* |  |
|  | TUGwalk | 137.88±54.75 | 0.029 |  | 154.66±64.02 | <0.001* |  | 145.99±61.74 | <0.001* |  | 112.29±44.27 | <0.001* |  |
|  | TUGstsi | 55.90±37.04 | <0.001* |  | 72.60±38.55 | <0.001* |  | 67.52±40.42 | <0.001* |  | 71.47±41.41 | <0.001* |  |
|  | acc | 158.85±66.10 | 1.000 |  | 170.58±69.55 | 1.000 |  | 173.86±76.51 | 1.000 |  | 118.81±47.33 | 1.000 |  |
|  | dec | 70.18±32.80 | <0.001* |  | 88.94±40.80 | <0.001* |  | 92.62±39.87 | <0.001* |  | 96.67±34.12 | 0.001* |  |
|  | stairA | 121.25±61.65 | 0.004* |  | 147.17±62.64 | <0.001* |  | 158.40±70.14 | 0.001* |  | 140.67±74.56 | 0.450 |  |
|  | stairD | 131.72±61.52 | 0.063 |  | 170.75±74.50 | 1.000 |  | 155.57±70.21 | <0.001* |  | 126.59±54.67 | 1.000 |  |
|  | stand | 41.53±18.49 | <0.001* |  | 45.20±17.02 | <0.001* |  | 52.06±19.91 | <0.001* |  | 60.57±24.41 | <0.001* |  |
| PTI (kPa.s) | self | 46.57±21.20 | - | <0.001* | 58.61±19.61 | - | <0.001* | 66.67±20.92 | - | <0.001* | 61.90±18.54 | - | <0.001* |
|  | TUGsist | 46.74±36.20 | 1.000 |  | 59.41±37.33 | 1.000 |  | 61.82±32.44 | 1.000 |  | 54.78±31.59 | 1.000 |  |
|  | TUGwalk | 48.06±22.91 | 1.000 |  | 57.77±20.93 | 1.000 |  | 62.21±18.97 | 0.043 |  | 59.49±18.71 | 0.599 |  |
|  | TUGstsi | 52.81±44.64 | 1.000 |  | 66.31±41.96 | 1.000 |  | 71.77±42.08 | 1.000 |  | 73.43±40.63 | 0.887 |  |
|  | acc | 49.92±26.26 | 1.000 |  | 59.66±19.60 | 1.000 |  | 66.10±20.49 | 1.000 |  | 63.12±18.82 | 1.000 |  |
|  | dec | 46.82±25.29 | 1.000 |  | 57.03±24.12 | 1.000 |  | 66.71±24.08 | 1.000 |  | 74.75±23.29 | 0.043 |  |
|  | stairA | 66.88±39.82 | 0.001* |  | 77.76±30.00 | <0.001* |  | 91.80±29.31 | <0.001* |  | 89.05±32.76 | <0.001* |  |
|  | stairD | 67.06±40.04 | 0.002* |  | 86.46±32.45 | <0.001* |  | 88.43±28.13 | <0.001* |  | 74.75±24.24 | <0.001* |  |
|  | stand | 33.59±17.06 | 0.001* |  | 35.75±15.40 | <0.001* |  | 43.41±18.25 | <0.001* |  | 52.09±23.46 | 0.179 |  |

\* = significantly different from walking at self-selected speed. Cut-off for p-values to be considered statistically significant was corrected for multiple testing with Holm-Bonferroni method. <sup>a</sup> = N can be less than 59 due to amputation. The F value shows if there was an overall significant difference between the means of different activities. (PPP = peak plantar pressure, PTI = pressure-time integral, self = walking at self-selected speed, TUG = Timed Up and Go test,; sist = sit-to-stand; walk = walking including turning; stsi = stand-to-sit,; acc = accelerating, dec = decelerating, stairA = stair ascending, stairD = stair descending, stand = standing)

## 4. Discussion

We investigated the effect of walking speed on plantar pressure measures, and compared plantar pressure measures of overground walking at self-selected speed with different weight-bearing activities of daily living, in people with diabetes at high risk of ulceration.

### 4.1. Speed conditions

Even at low walking speeds and with small differences between conditions (0.4, 0.6 and 0.8 m/s), PPP increased with increasing walking speed, while PTI decreased. This can be explained by an increase in ankle push-off at higher speeds, leading to an increase in ground reaction force and, with similar area of force application, higher PPP in all forefoot regions [31], while the step duration decreases, resulting in lower PTI [32]. The mean self-selected walking speed, 1.1 m/s, showed an even higher PPP and lower PTI compared to walking at 0.8 m/s. As such, PPP gives an overestimation and PTI an underestimation of the load on the foot in daily life when measured at self-selected speed under optimal conditions in a laboratory setting. With this inverse direction of PPP and PTI when walking speed changes, both may be valuable in describing the biomechanical load on the foot, even though the correlation between these parameters at self-selected speed is high [33].

### 4.2. Activity conditions

This is the first study that measured in-shoe plantar pressure during standing, and accelerating and decelerating phases of walking. Standing showed significantly lower PPP and PTI than walking at self-selected speed, due to a lack of ankle push-off. However, it should be considered that the time spent standing in daily life is twice as long as walking [24]. For accelerating and decelerating, ankle push-off increases during the first, while this decreases during the latter, explaining the similarity in PPP of accelerating and lower PPP of decelerating compared to walking at self-selected speed [31]. PTI was similar to walking at self-selected speed for both accelerating and decelerating, which is likely the result of a longer step duration for decelerating. During all other activities of daily living, PPP and PTI were similar or lower compared to walking at self-selected speed, in line with other studies in different populations [18–23]. The only exception was stair walking, which showed a higher PTI than walking at self-selected speed, also in line with previous studies [20,22], but it should be considered that the exposure to stair walking is limited in daily life compared to walking and standing [34]. In general, our results in people with diabetes at high risk of ulceration for whom plantar pressure measurements have proven clinical relevance, align with results of other populations.

### 4.3. Implications

#### 4.3.1. Cumulative plantar tissue stress

Although equipment for a valid continuous measurement of cumulative plantar tissue stress over multiple days is not yet available, it can be approximated by a model, for example by multiplying the PTI of mid-gait steps by the average number of strides per day [1]. However, this is a simplification of the cumulative stress experienced in daily life, because people walk at a wide range of speeds [8,9] and people undertake a variety of weight-bearing activities in daily life. This may explain the limited association found between current cumulative plantar tissue stress models and ulcer development [35] and ulcer healing [36], although investigated in two studies only. More comprehensive and therefore realistic cumulative plantar tissue stress models are needed, by including the type, frequency and duration of weight-bearing activities and walking speeds in daily life, along with the PTI of these activities and walking speeds [1]. This may help to better predict diabetes-related foot ulcer development and healing.

#### 4.3.2. Evaluation of offloading effect of therapeutic footwear

We showed that even at slow walking speeds and with differences of only 0.2 m/s between walking speeds, statistically significant changes in PPP and PTI were present. This implies for both research and clinical purposes that walking speed should be controlled for between repeated measurements in the evaluation of therapeutic footwear, for example when comparing different footwear conditions or footwear before and after adaptations [37]. Furthermore, insight in walking speeds in daily life in this high-risk population is needed. Currently, 200 kPa is used as a maximum threshold for in-shoe pressures to be protective against diabetes-related foot ulcers [38]; however, this is based on laboratory measurements during walking at self-selected speed. Because people walk slower in daily life, further development and personalization in this pressure threshold may be needed.

Our study indicates that PPP measured during self-selected speed in a laboratory is an overestimation of PPP to which people are exposed in daily life, because people walk faster in a laboratory than in daily life [8,9] and walking at self-selected speed showed the highest PPP when compared to other activities of daily living. The different components as assessed in the TUG may be a better representation of the load on the foot experienced in daily life, because the TUG consists of a combination of activities, and is executed at lower walking speeds. Future research should investigate whether plantar pressures measured during the TUG can be used to optimize footwear.

### 4.4. Strengths and limitations

A strength of our study was assessing the population with the highest risk of ulceration, where plantar pressure measurements have proven clinical relevance [2]. Furthermore, we measured plantar pressure during walking speeds and weight-bearing activities that are common in daily life. A limitation was that we did not randomize the order of conditions, which could result in influences of fatigue from the many conditions tested or a learning effect. However, this is unlikely, because the participants did not have to exert a maximum effort during measurements, they had sufficient rest between the measurements, all conditions are common in daily life, and plantar pressure is unlikely to be influenced by a learning effect. Another limitation may be that we measured plantar pressure while mimicking daily life conditions in a laboratory setting and not in people’s houses and environments. However, there are many variations in houses and environments, which would cause unwanted heterogeneity.

## 5. Conclusions

In people with diabetes at high risk of ulceration, we found that in-shoe PPP increases and PTI decreases significantly with increasing walking speed. PPP during activities of daily living was similar or lower compared to walking at self-selected speed. PTI was higher for stair walking, lower for standing and similar for other activities, compared to walking at self-selected speed. These differences should be considered when evaluating the biomechanical properties of therapeutic footwear, and when assessing cumulative plantar tissue stress.

## Author contributions

SAB, MP, JJvN, CMH and MGD Conceptualization; JJvN, CMH and MGD Data curation; CMH and MGD Formal analysis; JJvN and SAB Funding acquisition; JJvN, CMH and MGD Investigation; JJvN, CH, MGD, MP and SAB Methodology; JJvN and CMH Project administration; JJvN and CMH Resources; CMH and MGD Software; SAB, MP and JJvN Supervision; JJvN and CMH Validation; CMH and MGD Visualization; CMH and MGD Roles/Writing - original draft; SAB, MP and JJvN Writing - review & editing.

